# Diverse mutant selection windows shape spatial heterogeneity in evolving populations

**DOI:** 10.1101/2023.03.09.531899

**Authors:** Eshan S. King, Beck Pierce, Michael Hinczewski, Jacob G. Scott

## Abstract

Mutant selection windows (MSWs), the range of drug concentrations that select for drug-resistant mutants, have long been used as a model for predicting drug resistance and designing optimal dosing strategies in infectious disease. The canonical MSW model offers comparisons between two subtypes at a time: drug-sensitive and drug-resistant. In contrast, the fitness landscape model with *N* alleles, which maps genotype to fitness, allows comparisons between *N* genotypes simultaneously, but does not encode continuous drug response data. In clinical settings, there may be a wide range of drug concentrations selecting for a variety of genotypes. Therefore, there is a need for a more robust model of the pathogen response to therapy to predict resistance and design new therapeutic approaches. Fitness seascapes, which model genotype-by-environment interactions, permit multiple MSW comparisons simultaneously by encoding genotype-specific dose-response data. By comparing dose-response curves, one can visualize the range of drug concentrations where one genotype is selected over another. In this work, we show how *N*-allele fitness seascapes allow for *N ∗*2^*N−*1^ unique MSW comparisons. In spatial drug diffusion models, we demonstrate how fitness seascapes reveal spatially heterogeneous MSWs, extending the MSW model to more accurately reflect the selection fo drug resistant genotypes. Furthermore, we find that the spatial structure of MSWs shapes the evolution of drug resistance in an agent-based model. Our work highlights the importance and utility of considering dose-dependent fitness seascapes in evolutionary medicine.

**Author Summary:** Drug resistance in infectious disease and cancer is a major driver of mortality. While undergoing treatment, the population of cells in a tumor or infection may evolve the ability to grow despite the use of previously effective drugs. Researchers hypothesize that the spatial organization of these disease populations may contribute to drug resistance. In this work, we analyze how spatial gradients of drug concentration impact the evolution of drug resistance. We consider a decades-old model called the mutant selection window (MSW), which describes the drug concentration range that selects for drug-resistant cells. We show how extending this model with continuous dose-response data, which describes how different types of cells respond to drug, improves the ability of MSWs to predict evolution. This work helps us understand how the spatial organization of cells, such as the organization of blood vessels within a tumor, may promote drug resistance. In the future, we may use these methods to optimize drug dosing to prevent resistance or leverage known vulnerabilities of drug-resistant cells.

## Introduction

Drug resistance in cancer and infectious disease is governed by the unifying principles of evolution. Selection, which is integral to evolution, may be described by dose-response curves, which model growth rate as a function of drug concentration. Genotype-specific dose-response curves are ubiquitous across disease domains, including cancer and infectious disease. Dose-response curves can vary between different genotypes in multiple characteristics, such as their y-intercept (drug-free growth rate), *IC*_50_ (half-maximal inhibitory concentration), and shape^1–9^. Dose-response curves may also reveal fitness tradeoffs, where drug resistance imposes a fitness cost in the drug-free environment^10–12^. These diverse collections of dose-response curves among individual disease states give rise to varying degrees of selection when the drug concentration varies in time and space. Collections of genotype-specific dose response curves constitute fitness seascapes, which extend the fitness landscape model by mapping both genotype and environment (i.e. drug concentration) to fitness (**Box 1A**)^1–3,13–16^.

Box 1: Fitness seascape and mutant selection window terminology

#### Fitness seascapes

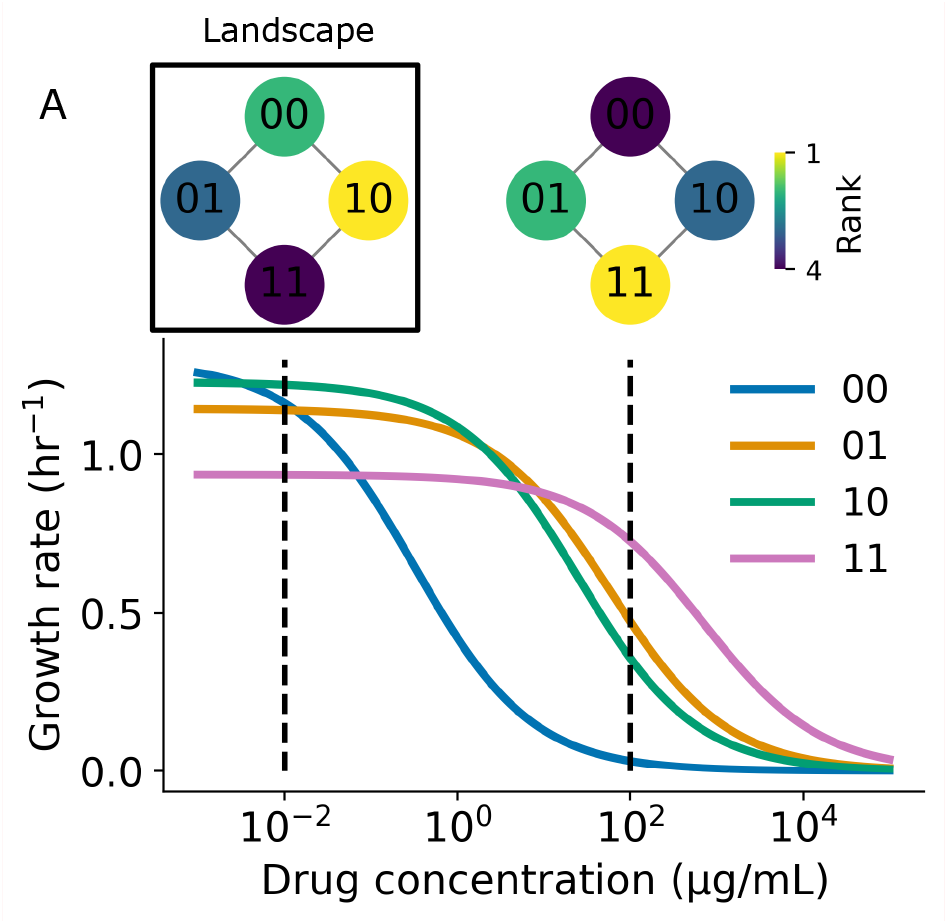

Fitness seascapes are extensions of the fitness landscape model that map both environment and genotype to fitness. For instance, while a fitness landscape may map genotype to minimum inhibitory concentration (MIC), a measure of drug resistance, a fitness seascape may map genotype and drug concentration to growth rate. Here, we model fitness seascapes as collections of genotype-specific dose-response curves. Previous usage of the term ‘fitness seascape’ has referred specifically to time-varying fitness landscapes. However, implicit in this use is that the environment shapes the fitness landscape, and it is the time-varying dynamics of the environment that result in the time-varying fitness landscape. In this work, we propose expanding the definition of fitness seascape to include any mapping from genotype and environment to fitness. Formally, we can define the relationship between fitness seascapes and fitness landscapes as

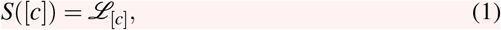

where *S*([*c*]) represents the fitness seascape as a function drug concentration and *L*_[*c*]_ represents the fitness landscape at a given drug concentration. In panel **A**, genotypes are modeled as binary strings of length 2, where each position in the string indicates the presence or absence of a particular point mutation. Each genotype is associated with a corresponding dose-response curve. Dose-response curves are modeled by:

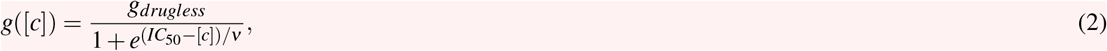

where *g*_*drugless*_ is the genotype-specific growth rate in the absence of drug, *IC*_50_ is the half-maximal inhibitory concentration, and *ν* is the Hill coefficient, which determines the steepness of the curve. The collection of genotype-specific dose response curves constitutes a fitness seascape, where fitness is a function of both genotype and drug concentration. Rank-order fitness landscapes that describe the relative fitness rank of each genotype at 10^*−*2^ and 10^2^ *μ*g/mL drug are shown inset. The fitness landscape changes as a function of drug concentration due to the dose-response curves associated with each genotype.

#### Mutant selection windows

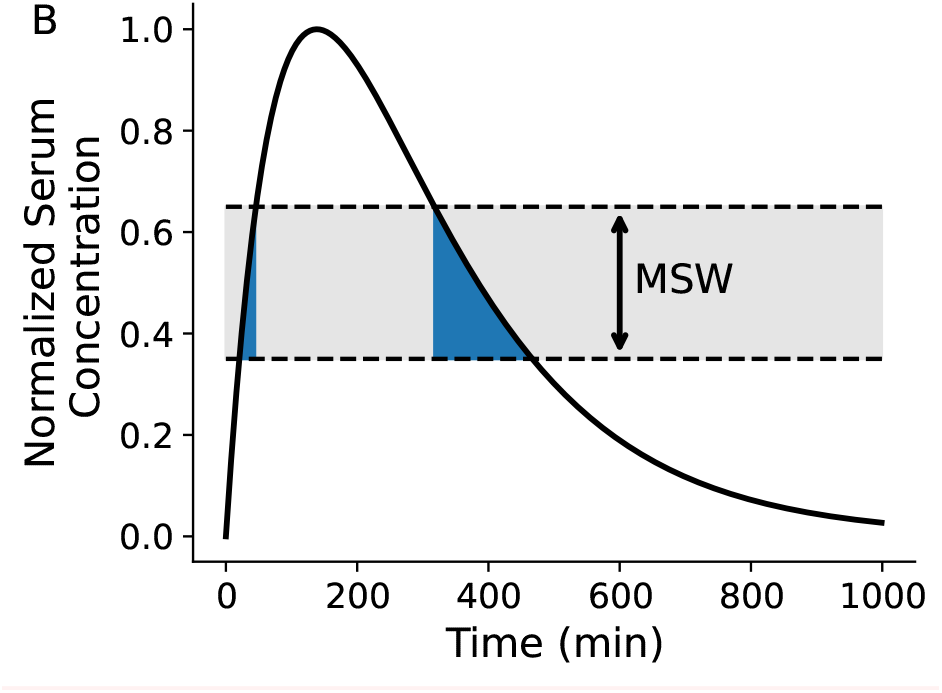

Mutant selection window refers to the range of antibiotic concentrations that select for a drug resistance mutant without fully suppressing growth^1,5,6,17–22^. MSWs are important to consider because they are thought to facilitate the evolution of antibiotic resistance. MSWs have been used to design dosing regimens that minimize the time that a patient’s serum drug concentration is within the MSW (*t*_*MSW*_)^21, 23^.

Panel **B** shows a simulated patient’s serum drug concentration following a single antibiotic dose. The MSW for this particular microbe is 0.35-0.65 in the normalized range. The time spent within the MSW, *t*_*MSW*_, is indicated by the blue shaded region. While the patient’s serum concentration is within this range, the drug resistant mutant is selected for without being fully suppressed.

The concept of the mutant selection window (MSW), the range of drug concentrations where the mutant growth rate is not fully suppressed and exceeds the wild-type growth rate, has been studied as a means of predicting evolution and optimizing drug regimens^1,5,6,17–22^. Under the MSW paradigm, drug regimens should be chosen that minimize the time a patient is subject to a drug concentration within the MSW (**Box 1B**)^22,24^. Das et al. have previously demonstrated that MSWs are intrinsically embedded in fitness seascapes^1^; by comparing dose-response curves in a fitness seascape, one can visualize the range of drug concentrations that selects for a drug resistant mutant. Previous work has shown how patient nonadherence and drug gradients caused by tissue compartmentalization confounds the use of MSWs in optimizing drug regimens, allowing for the emergence of drug resistant mutants^5,6,25,26^. Incorporating PK/PD models and MSWs may be crucial for translating evolutionary medicine to the clinic– for instance, Pan et al have demonstrated the presence of an MSW for *Pseudomonas aeruginosa* in an *in vivo* model, showing a correlation between sub-optimal PK/PD parameters and the emergence of drug resistance^27^.

MSWs traditionally offer comparisons between two genotypes or phenotypes at a time– i.e., between a drug susceptible reference genotype and a drug resistant genotype. By examining MSWs with the fitness seascape model, we may compare many genotypes simultaneously. For instance, in the binary landcsape model with *N* mutational sites, each genotype may be simultaneously compared to *N* adjacent genotypes (genotypes that differ by 1 mutation)^3,28–34^ Here, we expand on this idea and explore the MSW model through the lens of fitness seascapes. First, we illustrate how an *N*-allele fitness seascape allows for *N∗* 2^*N*^ MSW comparisons at a time. Then, we derive the steady-state drug concentration profile for drug diffusion from blood vessels in 1 and 2 dimensions, revealing the presence of heterogeneous MSWs. Finally, we explore how drug pharmacokinetics shapes MSW spatial heterogeneity and how MSW spatial structure impacts evolution using a 2-D agent-based model. While previous work has analyzed the importance of time spent in a MSW, we also consider the impact of space occupied by a MSW. We argue that both time and space occupied by a MSW may impact the emergence of resistance.

This work further explores the connection between mutant selection windows and fitness seascapes using realistic pharmacokinetic models. Our results highlight the utility of fitness seascapes in modeling evolution when drug concentration varies in time and space. Furthermore, because of the multiplicity of MSWs present in a fitness seascape, this work suggests that a higher-dimensional MSW model offered by fitness seascapes may be more powerful for predicting evolution in clinical settings, particularly when concerned with population heterogeneity.

## Results and Discussion

### Fitness seascapes embed mutant selection window data

We first investigated how multiple mutant selection windows are embedded in fitness seascapes, as demonstrated by Das et al.^1^. Here, we model genotypes as binary strings of length *N*, with a zero indicating no mutation and a 1 indicating a mutation in a given allelic position. *N* represents the total number of resistance-conferring mutations considered. These mutations can be thought of as either single nucleotide polymorphisms, single amino acid substitutions, or larger chromosomal changes. To illustrate this, we model a simple 2-allele fitness seascape by mapping a combinatorially complete set of genotypes (00, 01, 10, 11) to independent, randomly generated dose-response curves (**Box 1A**). Dose response curves differ in their IC_50_ and drug-free growth rates. Empirical fitness seascapes bearing a similar structure and demonstrating fitness tradeoffs have been reported by othersacross different kingdoms of life, providing a theoretical foundation for this approach^1–3^.

In an *N*-allele model, each genotype can be compared to *N* neighboring genotypes in the landscape (**Box 1A**). Here, ‘neighboring’ refers to genotypes that differ by one genetic change, or Hamming distance 1 (i.e., 00 and 01 are neighbors, but not 00 and 11). When calculating mutant selection windows, one first defines a wild type or ‘reference genotype’ to compare to the mutant. Given that each genotype in a fitness seascape can itself be thought of as the reference genotype and compared to its *N* neighbors, an *N*-allele fitness seascape embeds *N∗*2^*N*^ MSW comparisons. If one includes only *unique* MSW comparisons, i.e. the distinction between reference and comparison genotype is not meaningful, then this expression is *N* 2*∗*^*N−*1^. **Fig. 1B** shows the grid of all possible MSWs for a 2-allele seascape, shaded by the selection coefficient *s*_*i, j*_. The selection coefficient is defined as 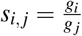, where *g*_*i*_ represents the growth of the more fit genotype and *g* _*j*_ represents the growth rate of the less fit genotype. Reference selection refers to the range of drug concentrations where the reference genotype has a higher fitness than the mutant. Similarly, mutant selection refers to the range of drug concentration where the mutant has a higher fitness. Net loss refers to the range of drug concentrations where the net replication rate of the reference and mutant genotypes are both less than 0.An example of how MSWs are calculated with dose-response curves is shown in **Fig. 1A**. Notably, the strength of selection blurs between the boundaries of the selection windows, further complicating the MSW paradigm (**Fig. 1B**). It may be useful to consider where selection is strongest within a MSW when designing dosing strategies.

**Figure 1.**
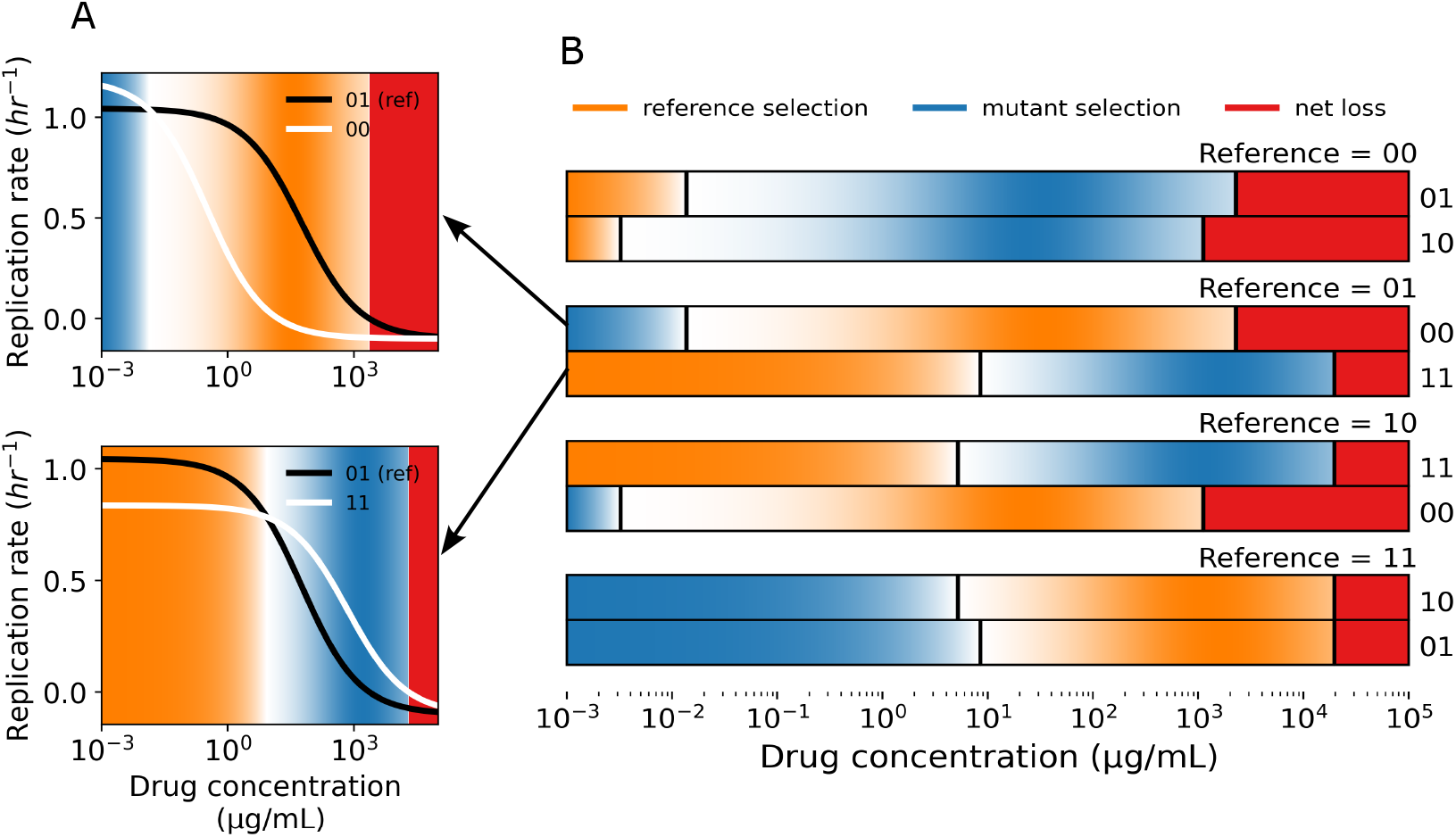
Fitness seascapes embed mutant selection window data. **A**) Example MSWs shown with the corresponding dose-response curves. The black line (01) is the dose-response curve for the reference genotype, while the white lines are considered the mutant genotypes. Orange corresponds to the reference selection window, blue to the mutant selection window, and red corresponds to the drug concentration that inhibits growth of both genotypes (net loss). **B**) All 8 (8 = *N∗* 2^*N*^, *N* = 2) MSW comparisons for a 2-allele fitness seascape. Each two row group represents a single reference genotype compared to its two neighboring genotypes. Reference selection, mutant selection, and net loss windows are calculated as a function of drug concentration and shown as colored rectangles. Selection windows are shaded by the normalized selection coefficient, *s*_*i, j*_, which is defined as the normalized ratio between the growth rates under comparison, 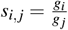, where *g*_*i*_ represents the growth rate of the genotype under selection and *g* _*j*_ represents the growth rate of the less fit genotype. All dose-response data is taken from the fitness seascape in **Box 1A**.

### Multiple mutant selection windows arise when drug concentration varies in space

We next sought to investigate MSWs in physiologically-relevant spatial models of drug drug diffusion in tissue. For the 1-dimensional (1D) case, we consider a location *x* at a time *t*. The drug concentration *u*(*x, t*) arising from a blood vessel source can be modeled with drug diffusion rate *D*, drug clearance rate *γ*, and the drug concentration at the tissue-blood vessel boundary *k*. We use a partial differential equation (**Eq. 3**) to find a steady state solution of drug diffusion:

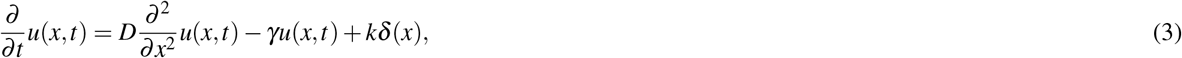

where *δ* (*x*) is the Kronecker delta function representing a blood vessel modeled as a point source. The 1D steady-state solution is of the form:

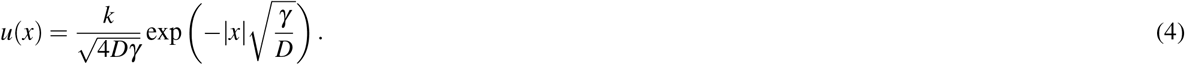

The clearance rate *γ* summarizes drug metabolism, consumption, and clearance into one parameter. We model a blood vessel source at *x* = 0 and compute drug concentration as a function of distance from the blood vessel. Using a similar approach to the time-varying case above, we used information provided by the simulated *N* = 2 fitness seascape to identify MSWs in space. Drug diffusion results in four different MSWs across the simulated 1D tissue patch (**Fig. 2A**). These results demonstrate that, given a constant supply of a drug source from a vessel in a tissue compartment, multiple mutant selection windows may exist simultaneously.

**Figure 2.**
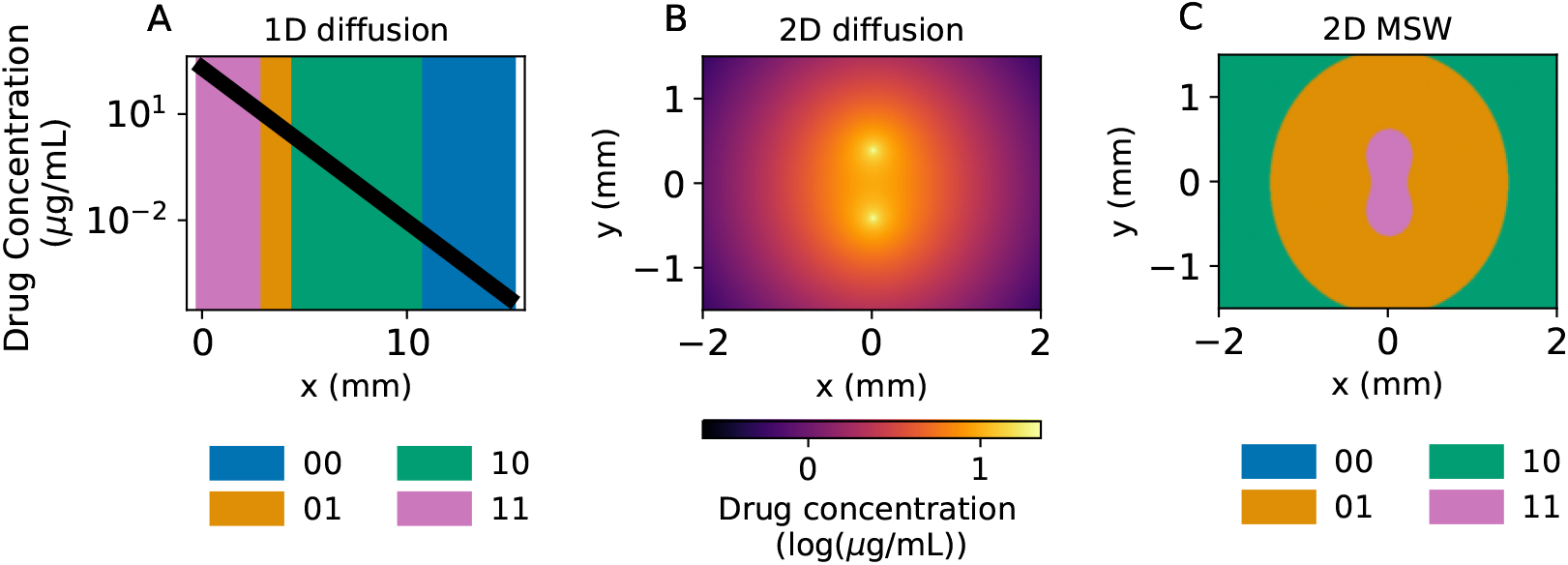
1- and 2-D diffusion models reveal diverse MSWs. **A)** Simulated 1-D drug diffusion in tissue from a blood vessel placed at x = 0. The black line represents the drug concentration as a function of distance *x* from the blood vessel, while the color represents the MSW at that region. **B)** 2-D steady state drug diffusion from two blood vessels placed at (0, −0.4) and (0, 0.4). **C)** Spatial MSWs corresponding to the steady-state diffusion in **B**. All dose-response data is based on the fitness seascape described in **Box 1A**.

To better understand spatial heterogeneity, we extended the model to 2 dimensions (2D) with a similar partial differential equation as before (see methods for derivation). The steady-state drug concentration is given by:

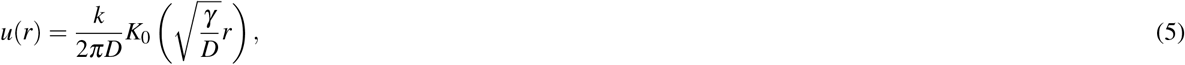

where *K*_0_( ) is a modified Bessel function of the second kind and *r* is the radial distance from the blood vessel source. We then considered the spatial effects of multiple blood vessels as drug sources in two dimensions, allowing for a more sophisticated representation of MSWs in tissue. For convenience, we set 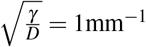. The distinction between 1 and 2 spatial dimensions is important, as the space occupied by a MSW in 2D scales by the distance from the blood vessel squared, versus linearly in 1D. The steady-state drug concentration profile is shown in **Fig. 2B**, with two blood vessels placed at (0,-0.4) and (0,0.4).

Identifying the MSWs as before reveals 3 distinct windows in this regime (**Fig. 2C**), with the 11 mutant window appearing to ‘bridge’ between the two blood vessels.

This 2D model offers a more complete picture of the MSWs functionally existing in an area around two blood vessels. These results suggest that including drug diffusion and pharmacodynamic effects may be important for simulating and predicting the evolution of drug resistance spatially. Introducing more blood vessels with arbitrary patterns would further complicate the drug diffusion pattern and the resulting MSWs. Together, these results complicate the notion of a single MSW driving the evolution of drug resistance. Instead, multiple MSWs may dictate evolution within a single population across time and space.

### Drug pharmacokinetics drive spatial heterogeneity

We next sought to understand how the spatial heterogeneity of MSWs may impact genetic heterogeneity in an evolving population. Using the Hybrid Automata Library (HAL)^35^, we implemented spatial agent-based simulations parameterized with the fitness seascape shown in **Box 1A**. In this work, we use the terms “agents” and “cells” interchangeably. We simulated evolution on a 100-by-100 grid with drug diffusion from two blood vessels placed at *x* = 50, *y* = 25 and *x* = 50, *y* = 75 (aligned vertically at the midline of the grid). Each simulation was initiated with a random sparsely populated lattice (initial density = 0.01 cells per grid position, on average) and an initial proportion of mutants of 0.1, meaning that each initial cell had a 0.1 probability of having a non-wild-type genotype (i.e. 01, 10, or 11). We also used HAL to simulate drug diffusion over time using the built-in partial differential equation (PDE) solver and the PDE grid functionality. Using the PDE solver, we studied how MSWs shift with different pharmacodynamic parameters, such as the drug elimination rate *γ*. The parameter *γ* may change depending on the drug under study, tissue type, and patient-specific drug metabolism. Example results are shown in **Fig. 3**, with each column in the figure corresponding to a value of *γ* increasing from left to right. We found that varying *γ* impacts the MSW pattern (**Fig. 3A** and **B**), with low elimination rate (*γ* = 10^*−*4^) selecting primarily for genotype 11 (pink), while a high elimination rate (*γ* = 0.5) selects primarily for genotype 00 (blue). Example simulations and summary data are shown in **Fig. 3C** and **D**.

**Figure 3.**
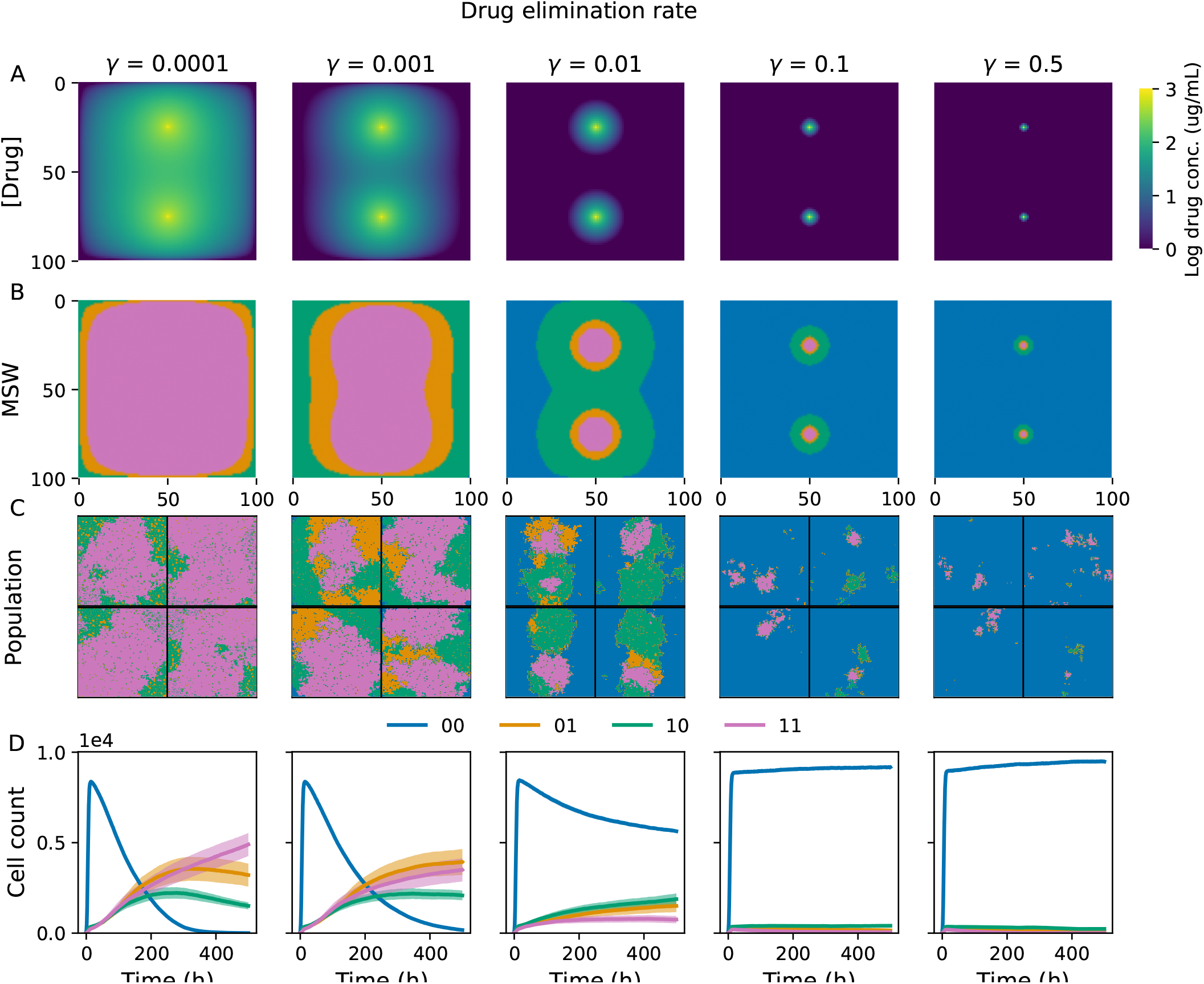
Varying drug elimination rate promotes MSW and population heterogeneity. Example results of simulations of cells evolving in a drug concentration gradient from two blood vessels. Each column corresponds to a different drug elimination rate (0 to 0.5, labeled at the top of each column). **A)** Final drug concentration profile at the end of the simulations. **B)** Mutant selection windows for the final drug concentration profiles. **C)** Example simulations. Each quadrant represents an individual simulation. Colors correspond to the cell genotype at each grid position. **D)** Averaged population counts from simulations for each condition. Colored lines correspond to the average number of cells of each genotype over time, while shading corresponds to the standard error estimate over time (*N* = 10 simulations per condition).

To better understand the utility of MSWs in predicting evolution, we next explored how the structure of MSWs, such as spatial heterogeneity, shapes evolution. First, we investigated the impact of MSW heterogeneity on the resulting population heterogeneity. We quantified MSW and population heterogeneity using Altieri entropy, which decomposes entropy into spatial mutual information and global residual entropy^36^. We found that MSW heterogeneity is strongly associated with population heterogeneity, with *R*^2^ = 0.84 (**Fig 4A**). The difference between the MSW entropy and mean population entropy at *γ* = 10^*−*4^ is likely due to the fact that the evolving population has not reached steady-state by the end of the simulation, as indicated by the changing cell counts in **Fig. 3D**. The initial condition for each simulation is a sparse grid of 90% wild-type (00), but the 11 mutant selection window takes up most of the grid by the end of the simulation.

**Figure 4.**
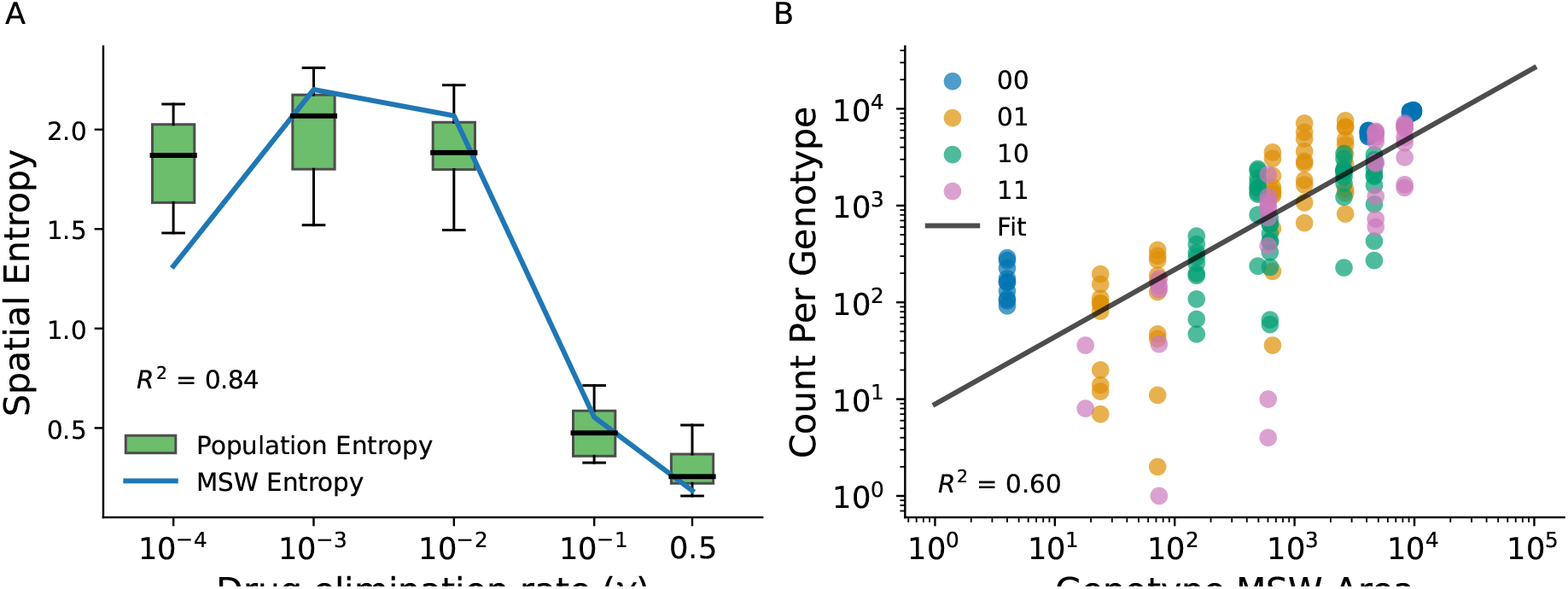
MSW structure drives population heterogeneity. **A)** Altieri spatial entropy of the population genotypes and MSWs as a function of drug elimination rate. *R*^2^ shown for the correlation between MSW entropy and population entropy. **B)** Correlation between the area occupied by an MSW and the number of cells of that genotype at the end of the simulation. Each point in the scatter plot corresponds to an individual genotype in a single simulation. *N* = 10 simulations per drug elimination rate *γ*.

Thus far, few studies have investigated the impact of space occupied by MSWs on the evolution of drug resistance – time spent within a MSW during a treatment regimen has been the primary concern when using MSWs to design optimal therapies. We found that the space occupied by a genotype’s mutant selection window is correlated with the number of cells of that genotype at the end of the simulation (**Fig 4B**), with *R*^2^ = 0.60. Taken together, these results suggest that 1) drug diffusion can drive MSW heterogeneity, 2) MSW spatial heterogeneity shapes population spatial heterogeneity, and 3) the area occupied by a mutant selection window is important to consider when studying the evolution of drug resistance.

### Sensitivity analysis of mutation rate, initial mutant proportion, and blood vessel geometry

We hypothesized that the predictive power of MSWs would depend on certain experimental parameters such as mutation rate and initial population heterogeneity, but would not depend environmental parameters such as blood vessel geometry. To test this, we performed a sensitivity analysis by varying initial mutant proportion, mutation supply, and blood vessel separation and quantifying the difference between the Altieri entropy of the MSWs and the population. Mutation supply is the mutation rate times the maximum population size (10^4^). Normalized entropy difference Δ_*entropy*_ was calculated as

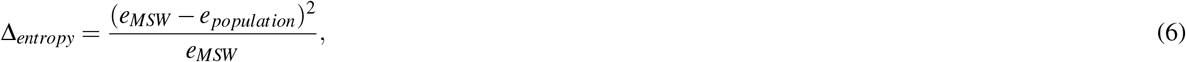

where *e*_*MSW*_ and *e*_*population*_ refer to the Altieri entropy of the MSWs and the population, respectively. Results of the sensitivity analysis are shown in **Fig. S1**. We found that Δ_*entropy*_ decreased with increasing mutation supply up to 100– beyond 100, drift begins to dominate over selection and the predictive power of MSWs decreases. The impact of initial mutant proportion on Δ_*entropy*_ was more prominent at lower mutation rates, with increasing initial mutant probability corresponding with lower Δ_*entropy*_. Interestingly, we found that Δ_*entropy*_ did not vary strongly with vessel separation (**Fig. S1B** and **Fig. S1C**), suggesting that the predictive capacity of MSWs is independent of blood vessel density.

## Discussion

In this work, we have investigated how MSW data is embedded in fitness seascapes and how numerous MSWs may exist when the selection pressure varies in time and space. In a simple model of drug diffusion from a blood vessel, we have illustrated how multiple MSWs may exist simultaneously in space. Furthermore, using a 2-D agent-based model of evolution, we showed how MSW structure shapes evolution of a population in a drug concentration gradient. Together, these results suggest that incorporating information from fitness seascapes may expand the power of MSW models to predict evolution.

Previous computational and experimental work has demonstrated that drug diffusion gradients and differential drug penetration, which permit heterogeneous MSWs, facilitate antibiotic resistance^25,^ . In addition, more complex blood vessel arrangements may confer additional MSW heterogeneity^38^. While the results presented here pertain to drug variation in space, variation in time is also a concern. Drug pharmacokinetic profiles, dosing regimes, and patient-specific nonadherence may cause the serum drug concentration to cross multiple mutant selection windows throughout the course of treatment. For instance, Nande and Hill, among others, have shown that drug absorption rate and patient nonadherence can impact the evolution of resistance^2,26^.

While previous applications of MSWs have focused on antibiotic resistance, we believe that the concepts explored here are also relevant to cancer, where drug resistance is a major driver of mortality. Cancer cells with different drug resistance mechanisms exhibit different dose-response curves, similar to microbes. Diffusion as a driver of MSW heterogeneity may be important in solid tumors, where nutrient and drug concentration gradients are thought to drive heterogeneity and contribute to drug resistance^38–42^. As the drug concentration profile is dependent on the tumor type, specific drug of interest, and vessel distribution, experiments accounting for these factors will be necessary for clinical translation of the theoretical concepts presented here. Previous work has demonstrated the use of radiolabeled drugs and immunofluorescence for quantifying drug diffusion *in vivo*^42^. Combining these techniques with microvasculature imaging technology such as super-resolution ultrasound imaging may enable fine-grained prediction of the drug concentration profile in an individual tumor^43^.

Our work is related to several other theoretical concepts across ecology, evolution, and medicine. For instance, fitness valley crossings, which permit adaptation despite there being a substantial fitness barrier, may be more likely in spatial gradients^44^; the boundary between mutant selection windows may promote genetic admixture, facilitating fitness valley crossings^45^. Furthermore, population heterogeneity discussed here may be thought of in the context of quasispecies theory^46,47^. Although quasispecies theory is commonly applied to viral dynamics, it may be more generally applied to any system with selection and genetic drift. A drug-dependent, spatial quasispecies theory may be comparable to the MSW model. While MSWs reflect selection only, quasispecies theory includes selection and drift and describes an equilibrium distribution of genotypes.

In the future, we may leverage fitness seascape data and the MSW analysis framework to more accurately predict the evolution of drug resistance. This may allow us to predict, or even control the trajectory of an evolving population and leverage known mechanisms of resistance^48–50^. Such an approach could reduce the total amount of drug used in the course of treatment and help mitigate the risk of drug resistance.

## Methods

### Diffusion from a point source with constant absorption in one dimension

We modeled the drug concentration gradient that a population of cells in tissue may experience as a point source with diffusion and constant absorption. The absorption rate encapsulates clearance, metabolism, and consumption of drug. Formally, we want to track the concentration *u*(*x, t*) on an infinite size domain, with diffusion described by diffusivity *D*, absorption at every position described by a rate *γ >* 0, and a source term that is a Kronecker delta function (*δ* (*x*)) with strength *k >* 0 at the origin *x* = 0. The diffusion equation then takes the form:

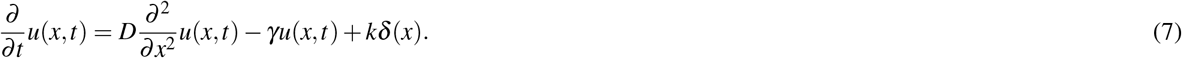

We assume we initially have zero concentration everywhere, so *u*(*x*, 0) = 0. To solve this, we will Fourier transform the above equation:

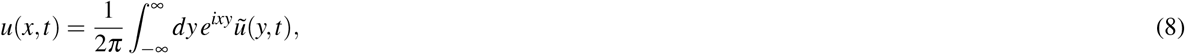

where *ũ* (*y, t*) is the Fourier transform of *u*(*x, t*). Now let us take temporal and spatial derivatives of Eq. (8), which we can then substitute into Eq. (7):

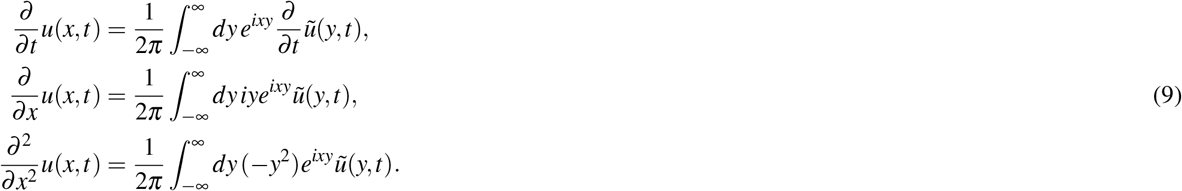

The final fact that we need is the Fourier transform of the Dirac delta function,

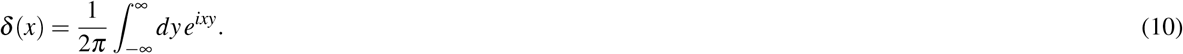

Substituting Eqs. (8)-(10) into Eq. (7), and collecting everything on one side under the same integral, we get:

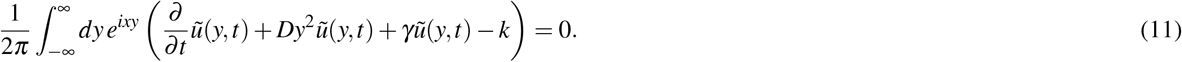

In order for Eq. (11) to be true for any *x*, the parenthetical terms have to equal to zero. This yields an ordinary, first-order differential equation for ũ (*y, t*):

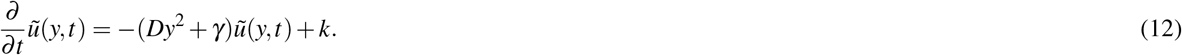

Using the initial condition *u*(*x*, 0) = 0, we find the solution for ũ (*y, t*):

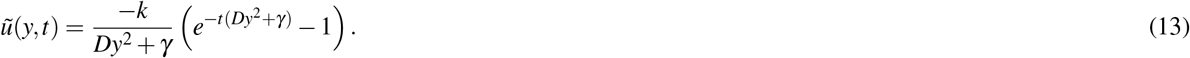

Substituting into Eq. (8) and rearranging we get our solution for *u*(*x, t*):

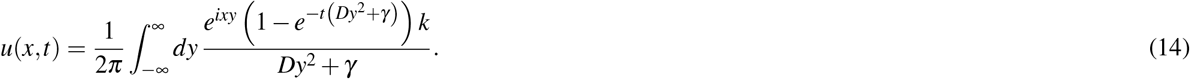

Taking the limit *t→* ∞, the concentration profile approaches a stationary distribution that is analytically solvable and simplifies the integration term to yield:

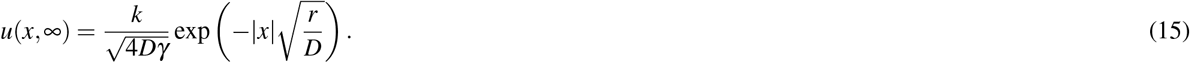

### Diffusion from a delta source with constant absorption in two dimensions

To model drug diffusion from a point source is two dimensions, we use a similar modeling technique as Eq. (7):

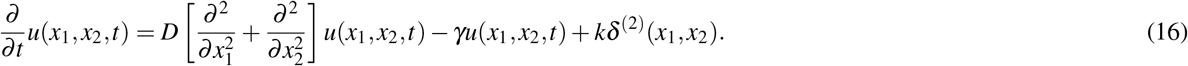

Here *δ* ^(2)^(*x*_1_, *x*_2_) is the 2D Dirac delta, *δ* ^(2)^(*x*_1_, *x*_2_) = *δ* (*x*_1_)*δ* (*x*_2_) which represents our drug source. We know by symmetry that the solution will only depend on the radial coordinate, *u*(*x*_1_, *x*_2_, *t*) = *u*(*r, t*), where 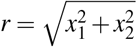. However, it is easier to proceed with the Fourier transform first in Cartesian coordinates, so we will delay the transformation to polar coordinates for now. The 2D Fourier transform is:

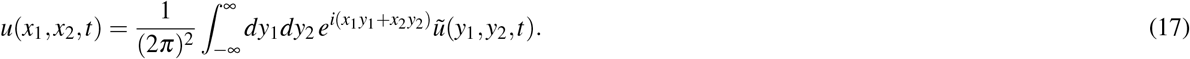

We calculate the temporal and spatial derivatives analogously to Eqs. (9)-(10), and the result is this 2D iteration of Eq. (11):

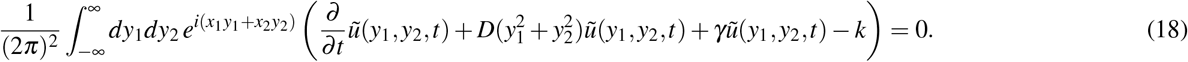

We can now convert our original and Fourier variables into polar coordinates: *x*_1_ = *r* cos *θ, x*_2_ = *r* sin *θ, y*_1_ = *ρ* cos *ψ, y*_2_ = *ρ* sin *ψ*. By symmetry we know that ũ (*y*_1_, *y*_2_, *t*) = ũ (*ρ, t*), where 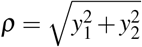 . Eq. (18) becomes:

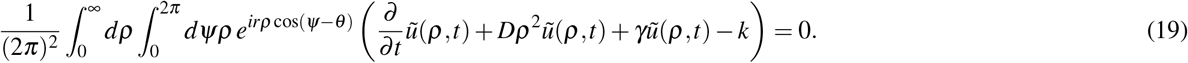

For this equation to be always true, the parenthetical terms must be zero, yielding:

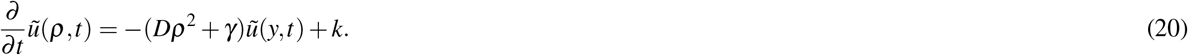

This has exactly the same form as Eq. (12) with *ρ* in place of *y*. Hence the solution mirrors Eq. (13):

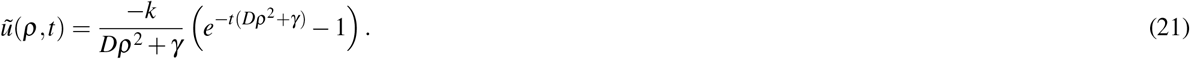

The solution *u*(*x*_1_, *x*_2_, *t*) = *u*(*r, θ, t*) in polar coordinates is then the inverse Fourier transform of the above:

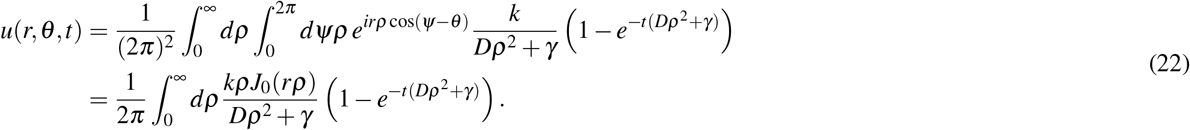

Here *J*_0_(*z*) is a Bessel function of the first kind. Note that the *θ* dependence has been integrated out, so *u*(*r, θ, t*) = *u*(*r, t*), as expected by symmetry. This integral cannot be computed analytically, but can be easily approximated numerically. To solve for the steady-state solution, we take the limit *t →* ∞:

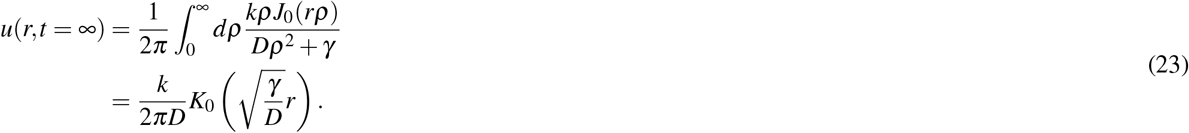

Here *K*_0_(*z*) is a modified Bessel function of the second kind. This function diverges at *r* = 0, which proves problematic for calculating concentrations near the blood vessel sources. Instead, we choose a blood vessel radius *r >* 0 and set the drug concentration within that radius equal to a constant maximum drug concentration. Thus, this analysis holds more strongly for analysis away from the immediate vicinity of the vessel.

To model drug diffusion from an arbitrary number of sources, we convolved the discretized version of Eq. (23) (*u*_*i, j*_(*t* = ∞)) with a 2D matrix of point sources (Δ_*i, j*_), where (*i, j*) represents position in the discretized 2D space:

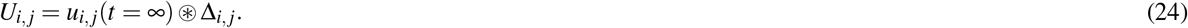

### Evolutionary simulations with HAL

We used HAL to implement on-lattice 2-dimensional agent-based simulations with drug diffusion. Each cell, or agent, was defined by its genotype, modeled by the binary strings 00, 01, 10, and 11, with each position in the string corresponding to resistance-conferring point mutation. These genotypes were assigned to the randomly generated dose-response curves shown in **Fig. 1A**, which determined the division probability of each cell as a function of drug concentration. At each time step, cells have the opportunity to divide with or without mutation, die, or do nothing. To divide, a cell must have at least one adjacent grid space unoccupied. When mutating, cells can change genotype to any of the two “adjacent” genotypes, meaning genotypes which differ by 1 position in the binary string (i.e., genotype 10 can mutate to 11 or 00 with equal probability). A lattice of size 100×100 was used for all simulations. For the simulations used in **Fig. 3** and **Fig. 4**, blood vessels were placed at positions x = 50, y = 25 and x = 50, y = 75. Diffusion was modeled using the PDE solver functionality in HAL, with the blood vessels supplying a constant drug concentration. Each point in the lattice absorbed drug at a variable rate *γ*, and the boundary was set to drug concentration of 0. Parameters used in the simulations for **Fig. 3** and **4** are shown in **Table 1**.

**Table 1.** Simulation parameters.

| Parameter | Value |
| --- | --- |
| Mutation rate | $10^{-4} \text{ bp}^{-1}$ |
| Initial mutant probability | 0.1 |
| Initial grid density | 0.01 |
| Blood vessel drug concentration | $10^3 \mu\text{g/mL}$ |
| Diffusion rate | 0.1 |
| Time steps | 500 hr |

For the sensitivity analyses, mutation rate and initial mutant probability were varied between 0 and 10^*−*1^ and blood vessel spatial separation was varied from 0 to 80 grid points (**Fig. S1**).

## Data availability

All code and data required for simulations, data analysis, and figure reproduction can be found at https://github.com/eshanking/msw_analysis/.

## Acknowledgements

JGS and ESK were supported by NIH 5R37CA244613-04 (https://www.cancer.gov/). ESK was supported by NIH 3T32GM007250-46S1 (https://www.nigms.nih.gov/). JGS was supported by American Cancer Society Research Scholar Grant RSG-20-096-01 (https://www.cancer.org/). The funders had no role in study design, data collection and analysis, decision to publish, or preparation of the manuscript. The authors also thank Paulameena Shultes for her insightful editorial feedback.

## Contributions

ESK: Conceptualization, formal analysis, investigation, methodology, software, visualization, writing – original draft preparation. BP: Formal analysis, investigation, software, visualization, writing – original draft preparation. MH: Formal analysis, investigation, methodology, supervision, writing – review and editing. JGS: Conceptualization, funding acquisition, resources, supervision, writing – review and editing.

## Supplemental Materials

### Sensitivity analysis of mutation rate, initial mutant probability, and blood vessel separation

We hypothesized that the predictive utility of mutant selection windows would depend on certain experimental parameters, such as mutation rate and initial mutant probability, but not environmental parameters such as blood vessel geometry. We performed a sensitivity analysis by quantifying the difference between the MSW and population Altieri spatial entropy as a function of mutation supply, initial mutant probability, and blood vessel spatial separation. Mutation supply is defined as the mutation rate multiplied by the maximum population size (10^4^). Sensitivity analyses were performed as an exhaustive sweep over all combinations of two parameters at a time. We first varied mutation supply and initial mutant probability, finding that the entropy difference decreased as mutation supply increased from 0 to 100 (**Fig. S1A**). For a mutation supply of 1000 (corresponding to a mutation rate of 0.1), drift begins to dominate over selection, limiting the predictive power of MSWs. Similarly, entropy difference decreased as initial mutant probability increased for mutation supply less than 100. However, this trend was ameliorated for a mutation supply of 100 and 1000, suggesting that drift dominates over initial population heterogeneity in these regimes. When we varied initial mutant probability and blood vessel separation, we once again found that the entropy difference decreased as initial mutant probability increased (**Fig. S1B** and **D**). However, there was not a significant trend associated with blood vessel separation (**Fig. S1C**). Example MSWs and population distributions for the blood vessel separation experiment are shown in **Fig. S2**.

**Figure S1. Sensitivity analysis reveals model robustness to vessel geometry.** A) Heatmap of mutation supply (mutation rate*population supply) versus initial mutant probability. Normalized entropy difference is the squared difference between the Altieri entropy of the MSW map and the Altieri entropy of the final population distribution, normalized by the MSW entropy (Eq 6). B) Heatmap of vessel separation (lattice points between the two blood vessels) versus initial mutant probability. C) Normalized difference in Altieri entropy versus blood vessel separation, averaged across initial mutant probabilities in panel B. D) Normalized difference in Altieri entropy versus initial mutant probability, averaged across the blood vessel separation values in panel B.

**Figure S2. Example MSWs and population distributions for vessel separation experiments.** Each column corresponds to a different blood vessel separation distance labeled at the top of each column. Distance is in units of lattice points. Drug elimination rate *γ* = 0.01, mutation rate = 0.0001, and initial mutant probability = 0.01.

