## Supplementary figures and images for "Diverse mutant selection windows shape spatial heterogeneity in evolving populations"

### Supplemental Figure 1

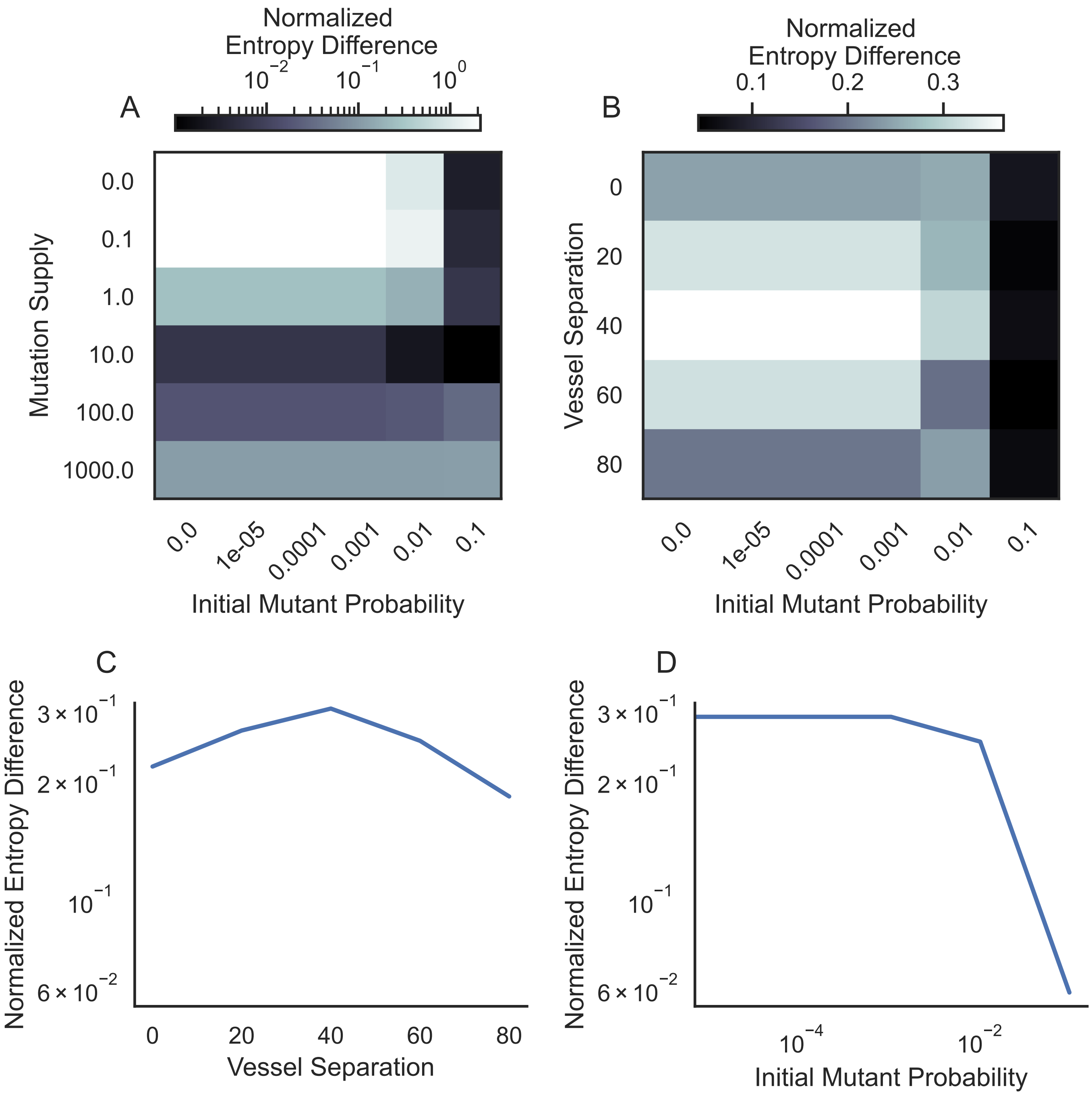

### Supplemental Figure 2

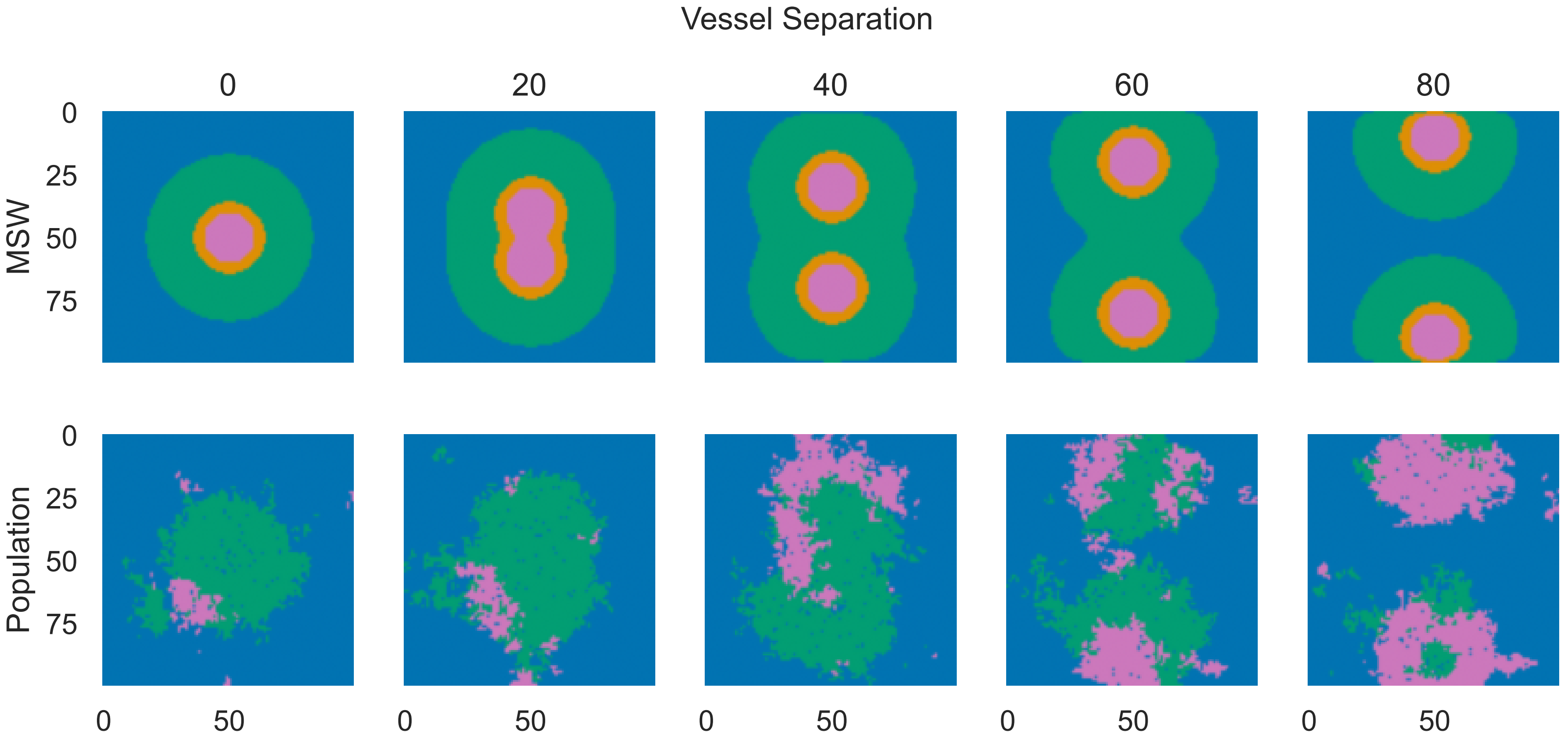
